## Supplementary Material for "#GotGlycans: Role of N343 Glycosylation on the SARS-CoV-2 S RBD Structure and Co-Receptor Binding Across Variants of Concern"

**Material and Methods**

**MD Simulations.** The RBD simulation systems were constructed from residues R327 to N540 of the SARS-CoV-2 S glycoprotein. The starting structure for WHu-1 MD1 was from the open RBD obtained from the simulation of the S ectodomain(Casalino et al., 2020), and the WHu-1 MD2 starting structure from PDB 6M0J. The alpha (B.1.1.7) starting structure was obtained from PDB 6M0J, with the N501Y mutation introduced with the mutagenesis tool in pymol (<https://pymol.org/2/>). The beta starting structures were from PDB 7LYN. All delta starting structures (MD1 and MD2, glycosylated and non-glycosylated) were obtained from PDB 7V7Q, with MD1 and MD2 productions started from different velocities. The omicron starting structures for MD1 was from PDB 7WVN, and the starting structure for MD2 from PDB 7QO7. Simulations of BA.2.86 were performed from PDB 7WVN, with sequence mutations introduced with the mutagenesis tool in pymol.

The charged N- and C-terminal residues were neutralised by capping with acetyl (ACE) and N-methylamide (NME) groups, respectively. The RBD was glycosylated with FA2G2 glycans at N331 and at N343. The structure of the (GlyTouCan-ID G00998NI) N-glycan was sourced from the GlycoShape glycan structure database (GDB), and was linked to the RBD using the Re-Glyco, a glycoprotein builder tool developed for GlycoShape (<https://glycoshape.org>). The systems were solvated in a water box with a minimum distance of 12 Å, and ions were added to neutralise any system charges to a total concentration of 200 mM NaCl.

All simulations were performed using AMBER18(D.A. Case, I.Y. Ben-Shalom, S.R. Brozell, D.S. Cerutti, T.E. Cheatham, III, V.W.D. Cruzeiro, T.A. Darden, R.E. Duke, D. Ghoreishi, M.K. Gilson, H. Gohlke, A.W. Goetz, D. Greene, R Harris, N. Homeyer, Y. Huang, S. Izadi, A. Kovalenko, T. Kurtzman, T.S. Lee, S. LeGrand, P. Li, C. Lin, J. Liu, T. Luchko, R. Luo, D.J. Mermelstein, K.M. Merz, Y. Miao, G. Monard, C. Nguyen, H. Nguyen, I. Omelyan, A. Onufriev, F. Pan, R. Qi, D.R. Roe, A. Roitberg, C. Sagui, S. Schott-Verdugo, J. Shen, C.L. Simmerling, J. Smith, R. SalomonFerrer, J. Swails, R.C. Walker, J. Wang, H. Wei, R.M. Wolf, X. Wu, L. Xiao, D.M. York and P.A. Kollman, 2018) on resources provided by the Irish Centre for High-End Computing (ICHEC). The AMBER 14SB force field(Maier et al., 2015) was used to model proteins and ions, the GLYCAM06j-1(Kirschner et al., 2008) force field was used to model glycans, and the TIP3P water model was used to model solvent molecules(Jorgensen et al., 1983).

The energy of the system was minimised in 500,000 steps using the steepest descent algorithm, with all heavy atoms of the protein and glycan restrained with a potential weight of 5 kcal.mol^-1^.Å^-2^. The system was then equilibrated in the NVT ensemble, with the system gradually heated from 0 to 100 K, and then from 100 K to 300 K. The system was then equilibrated in the NPT ensemble to maintain the pressure at 1 bar. Following this, most restraints were then removed, with restraints of a magnitude of 5 kcal.mol^-1^.Å^-2^ remaining in place on the caps, R327-N334, and G526-N540. This is because these regions of the RBD would be connected to the other domains of the spoke glycoprotein, and so would be structurally constrained in vivo. The system was then equilibrated for a further 100 ns, before production runs of ~2 μs, as summarised in **Table S.1**.

The temperature was maintained at 300 K using Langevin dynamics with a collision frequency of 1 ps^-1^, and the pressure was maintained at 1 bar using isotropic position scaling with a Berendsen barostat and a pressure relaxation time of 2 ps. Periodic boundary conditions were used throughout the simulations. The Van der Waals interactions were truncated at 11 Å and Particle Mesh Ewald (PME) was used to treat long range electrostatics with B-spline interpolation of order 4. The SHAKE algorithm was used to constrain all bonds containing hydrogen atoms and to allow the use of a 2 fs time step for all simulations.

**Table S.1** Molecular Dynamics (MD) sampling methods and corresponding times in the production phase of the simulations. MD1 and MD2 indicate trajectories collected through conventional (deterministic) sampling. GaMD indicates trajectories obtained through Gaussian accelerated MD sampling. Further details on the methodology and starting structures are in the text.

|  | **N343-FA2G2 (μs)** | | | **N343 NoGly (μs)** | | |
| --- | --- | --- | --- | --- | --- | --- |
| **Variant** | **MD1** | **MD2** | **GaMD** | **MD1** | **MD2** | **GaMD** |
| WHu-1 | 2.5 | 2.0 | 2.0 | 3.0 | 2.0 | 2.0 |
| Alpha (B.1.1.7) | 2.0 | - | - | - | 2.0 | - |
| Beta (B.1.351) | 2.0 | - | - | - | 2.0 | - |
| Delta (B.1.617.2) | 2.0 | 2.0 | 2.0 | 3.0 | - | 2.0 |
| Omicron (BA.1) | 2.0 | 2.0 | 2.0 | 2.0 | 2.0 | 2.0 |
| Omicron (BA.2.86) | 2.0 | - | - | - | - | - |
| **Total (46.5)** | **12.5** | **6.0** | **6.0** | **8.0** | **8.0** | **6.0** |

Additionally, Gaussian-accelerated MD (GaMD) simulations(Miao et al., 2015; Wang et al., 2021) were run using the GaMD module in AMBER18(D.A. Case, I.Y. Ben-Shalom, S.R. Brozell, D.S. Cerutti, T.E. Cheatham, III, V.W.D. Cruzeiro, T.A. Darden, R.E. Duke, D. Ghoreishi, M.K. Gilson, H. Gohlke, A.W. Goetz, D. Greene, R Harris, N. Homeyer, Y. Huang, S. Izadi, A. Kovalenko, T. Kurtzman, T.S. Lee, S. LeGrand, P. Li, C. Lin, J. Liu, T. Luchko, R. Luo, D.J. Mermelstein, K.M. Merz, Y. Miao, G. Monard, C. Nguyen, H. Nguyen, I. Omelyan, A. Onufriev, F. Pan, R. Qi, D.R. Roe, A. Roitberg, C. Sagui, S. Schott-Verdugo, J. Shen, C.L. Simmerling, J. Smith, R. SalomonFerrer, J. Swails, R.C. Walker, J. Wang, H. Wei, R.M. Wolf, X. Wu, L. Xiao, D.M. York and P.A. Kollman, 2018). GaMD simulations were performed for glycosylated and non-glycosylated RBD systems of the Whu-1, delta, and omicron variants. For these GaMD simulations, the starting structure for the Whu-1 variant was obtained from the simulation of the S ectodomain(Casalino et al., 2020), and the starting structures for the delta and omicron variants from PDBs 7V7Q and 7WVN, respectively. In brief, a 2 ns short conventional MD simulation was used to collect the required potential statistics for calculating GaMD acceleration parameters. This was followed by a 50 ns equilibration after adding the boost potential, and finally a 2 μs GaMD production simulation. The average and standard deviation of the system potential energies were calculated every 0.4 ns. GaMD simulations were run at the “dual-boost” level by setting the reference energy to the lower bound. One boost potential was applied to the dihedral energetic term, and the other to the total potential energetic term. The upper limit of the boost potential standard deviation was set to 6 kcal.mol^-1^ for both the dihedral and total potential energetic terms. The same temperature and pressure parameters were used in the conventional MD simulations.

**Structure of the RBD-GM1o Complex.** The equilibrium structure of the unbound RBD was obtained from MD simulations of the SARS-CoV-2 spike (S) WHu-1 ectodomain (Harbison et al., 2022) and was screened for binding to the GM1 tetrasaccharide (GM1o)(Nguyen et al., 2021). The GM1o structure was built with the GLYCAM Carbohydrate Builder and equilibrated by MD simulations in bulk water separately from the RBD. The same GM1o structure is deposited in the GlycoShape GDB (<https://glycoshape.org>) where it can be downloaded. As docking did not produce any convincing binding poses, none of which proved to be stable when tested by MD simulations, we used the conformation of the N370 N-glycan bound across the RBD(Harbison et al., 2022) as a guideline for the generation of a set of potential RBD/GM1o complexes. The stability of all promising conformations produced this way was tested by extensive sampling through MD simulations, started with a restrained equilibration phase (100-500 ns), where the protein was equilibrated around the bound (fixed) GM1o, followed by unrestrained MD. The complex shown in **Figure 1**, **panel d)** was obtained after numerous binding and unbinding events that occurred during the unrestrained MD, and it remained stable for 700 ns. As an interesting note, the position of the Neu5Ac in the bound GM1o corresponds to the position that a terminal sialic acid in N370 would have when bound to the RBD.

**Assignment of glycan compositions of RBD glycoforms**

For each RBD VOCs, the data were analysed using the measured molecular weights (MWs) of intact protonated WT, Alpha, Beta, Delta and Omicron RBD (WTx, Ax, Bx, Dx, and Ox respectively, **Tables S.2-6**). The MW of deglycosylated RBD was calculated based on the elemental composition corresponding to amino acid sequence (EG^319^RVQP…VN^541^FS, UniProt number P0DTC2) and any amino acid substitutions plus FLAG tag (SGDYKDDDDKG) and hexa histidine tag (HHHHHHG) and four disulfide bonds. The MWs of non-glycosylated RBD WT, Alpha, Beta, Delta and Omicron are 27,352.6 Da, 27,401.6 Da, 27,386.6 Da, 27,422.7 Da and 27,613.9 Da, respectively. Possible glycan compositions were simulated for the numbers of N-acetylhexosamines (N: HexNAc, N-acetylgalactosamine and N-acetylglucosamine), hexoses (H: Hex, glucose and galactose), fucoses (F: Fuc) and N-acetylneuraminic acids (S: Neu5Ac). Possible values of N, H, F and S were calculated by considering reported *N*- and *O*-glycans. Possible MWs of RBD glycoforms were then calculated from the sum of aforementioned MW of RBD and MWs of glycan residues from each possible H_N_F_S combination.

**ESI-MS affinity measurements**

The affinities of glycan ligands for RBD were measured by the direct ESI-MS binding assay(Kitova et al., 2012). For a monovalent protein–ligand (PL) interaction (eq. 1), the dissociation constant (*K*_d_; eq. 2) can be calculated from the ratio (*R*) of total abundances of L-bound (*Ab*(PL)) to free P ions (*Ab*(P), eq. S3) measured by ESI-MS

P+L ⇌ PL	(S1)

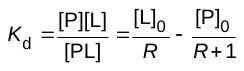
 (S2)

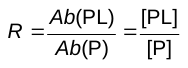
(S3)

where [P]_0_ and [L]_0_ are initial concentrations of P and L, respectively. The abundance ratio *R* measured by ESI-MS is taken to be equal to the equilibrium concentration ratio in the solution. Because RBD consists of multiple species with distinct glycan compositions, and glycosylation can, in principle, influence binding, the affinities for L binding to individual RBD species (eqs. S4a,b) were determined.

P_1_ + L ⇌ P_1_L	(S4a)

P_x_ + L ⇌ P_x_L	(S4b)

The corresponding equations of mass balance equations are

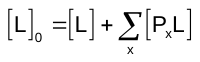
(S5a)

[P_1_]_0_ = [P_1_] + [P_1_L]	(S5b)

[P_2_]_0_ = [P_2_] + [P_2_L]	(S5c)

[P_x_]_0_ = [P_x_] + [P_x_L]	(S5d)

where [P_x_]_0_ is the initial concentration of a given P_x_ species. Initial concentrations of individual RBD species were estimated from their relative abundances measured by ESI-MS, assuming uniform response factors (eq. S6).

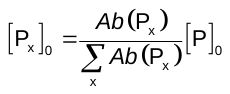
(S6)

The affinity of a given P_x_ species (*K*_dx_) was calculated from eq. S7

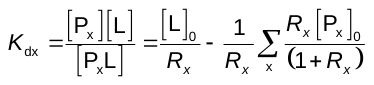
(S7)

where *R_x_* is the total abundance ratio of ligand-bound and free P*_x_* ions (eq. S8).

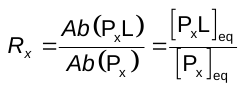
 (S8)

In some instances, signal for one ligand-bound P_x_ complex overlapped with signal for another (free) P_x_ species. Spectral overlap was corrected for by considering the abundance ratio of the two P_x_ species (for example P_x_ and P_x+1_) in the absence of L (*r*; eq. S9) and assuming that P_x_ and P_x+1_ exhibit identical affinities for L.

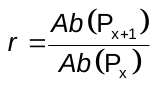
 (S9)

The corresponding *r* value was then used to calculate the true *R*_x_ value (*R_x,corr_*; eq. S10) corrected for spectral overlap,

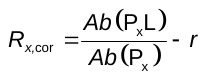
 (S10)

which was used in eq. S7 to calculate *K*_dx_.

**Additional Tables**

**Table S.2.** Affinities of the oligosaccharides of GM1 and GM2 (GM1_os_ and GM2_os_, respectively) for WHu-1, Alpha, Beta, Delta and Omicron RBD and endoH-treated WHu-1 RBD and endoF3-treated WHu-1 and Delta RBD measured by ESI-MS for aqueous ammonium acetate (100 mM, pH 7.4) solutions containing a given RBD (5  μM) and glycan (three different initial concentrations ranging from 10 μM to 150 μM). Data represent mean ± s.d.; n = 3 independent experiments for each glycan concentration.

| **Variant** | **GM1_os_ *K*_d_ (mM)** | **GM2_os_ *K*_d_ (mM)** |
| --- | --- | --- |
| WHu-1 RBD (sample 1; HEK293-a) | 0.16 土 0.04 | 0.17 土 0.02 |
| WHu-1 RBD (sample 2; HEK293-a) | 0.18 土 0.01 | 0.22 土 0.03 |
| WHu-1 RBD (sample 3; HEK293-b) | 0.07 土 0.01 | 0.09 土 0.02 |
| endoF3-treated WHu-1 RBD (HEK293-b) | 3.6 土 0.7 | 5.7 土 0.6 |
| alpha (HEK293-b) | 0.15 土 0.03 | 0.16 土 0.03 |
| beta (HEK293-b) | 1.3 土 0.4 | 1.10 土 0.2 |
| delta (HEK293-b) | 0.11 土 0.01 | 0.08 土 0.02 |
| endoF3-treated delta (HEK293-b) | 0.12 土 0.01 | 0.06 土 0.01 |
| omicron (HEK293-b) | 1.1 土 0.2 | 0.70 土 0.16 |

**Table S.3.** Summary of molecular weights (MWs) of WHu-1 RBD glycoforms (WTx) with respective relative abundances identified by ESI-MS and putative H_N_F_S combinations.

| RBD | Measured  MW (Da) | Relative abundance | H_N_F_S combination | Theoretical MW (Da) | Mass difference (Da) |
| --- | --- | --- | --- | --- | --- |
| WT1 | 31342.4 | 25.5 | 9_11_2_0 | 31345.1 | 5.3 |
| WT2 | 31402.4 | 2.6 | 11_9_1_1 | 31408.0 | 2.4 |
| WT3 | 31505.2 | 42.7 | 9_9_0_3 | 31520.1 | 6.9 |
| WT4 | 31527.0 | 34.3 | 10_9_1_2 | 31537.1 | 2.1 |
| WT5 | 31609.4 | 19.0 | 11_10_3_0 | 31612.1 | 5.3 |
| WT6 | 31668.8 | 48.8 | 10_9_0_3 | 31682.1 | 5.4 |
| WT7 | 31690.2 | 56.2 | 11_9_1_2 | 31699.1 | 1.0 |
| WT8 | 31711.4 | 17.8 | 9_10_0_3 | 31723.2 | 3.7 |
| WT9 | 31771.1 | 43.9 | 12_10_3_0 | 31774.2 | 4.9 |
| WT10 | 31794.0 | 65.6 | 10_11_0_2 | 31797.2 | 4.8 |
| WT11 | 31816.5 | 26.5 | 10_9_1_3 | 31828.2 | 3.7 |
| WT12 | 31835.3 | 30.6 | 11_9_2_2 | 31845.2 | 1.9 |
| WT13 | 31854.4 | 44.6 | 9_10_1_3 | 31869.2 | 6.8 |
| WT14 | 31875.2 | 47.5 | 10_10_2_2 | 31886.2 | 3.1 |
| WT15 | 31896.4 | 35.8 | 11_10_1_2 | 31902.2 | 2.2 |
| WT16 | 31935.8 | 5.6 | 10_11_1_2 | 31943.3 | 0.6 |
| WT17 | 31957.8 | 85.0 | 11_11_0_2 | 31959.2 | 6.6 |
| WT18 | 31979.7 | 98.1 | 13_11_0_1 | 31992.3 | 4.6 |
| WT19 | 31001.2 | 61.3 | 9_10_2_3 | 32015.3 | 6.1 |
| WT20 | 32038.7 | 17.3 | 11_10_0_3 | 32047.3 | 0.6 |
| WT21 | 32060.1 | 55.7 | 12_10_1_2 | 32064.3 | 3.9 |
| WT22 | 32083.3 | 55.9 | 10_11_0_3 | 32088.3 | 3.0 |
| WT23 | 32105.5 | 1.1 | 12_11_0_2 | 32121.3 | 7.8 |
| WT24 | 32125.1 | 12.6 | 13_11_1_1 | 32138.3 | 5.2 |
| WT25 | 32144.3 | 72.6 | 10_12_1_2 | 32146.3 | 5.9 |
| WT26 | 32165.0 | 63.5 | 12_12_1_1 | 32179.3 | 6.4 |
| WT27 | 32185.7 | 39.8 | 11_10_1_3 | 32193.3 | 0.4 |
| WT28 | 32247.8 | 88.4 | 11_11_0_3 | 32250.3 | 5.5 |
| WT29 | 32270.3 | 100.0 | 13_11_0_2 | 32283.4 | 5.1 |
| WT30 | 32291.8 | 62.4 | 12_14_1_0 | 32294.4 | 5.4 |
| WT31 | 32348.8 | 58.7 | 12_10_1_3 | 32355.4 | 1.4 |
| WT32 | 32371.5 | 46.6 | 10_11_0_4 | 32379.4 | 0.1 |
| WT33 | 32434.4 | 69.7 | 10_12_1_3 | 32437.4 | 5.0 |
| WT34 | 32456.1 | 76.4 | 12_12_1_2 | 32470.4 | 6.3 |
| WT35 | 32477.0 | 43.6 | 11_10_1_4 | 32484.4 | 0.6 |
| WT36 | 32538.2 | 78.0 | 11_11_0_4 | 32541.4 | 4.8 |
| WT37 | 32560.9 | 79.6 | 13_11_0_3 | 32574.4 | 5.6 |
| WT38 | 32583.0 | 49.5 | 12_14_1_1 | 32585.5 | 5.5 |
| WT39 | 32615.2 | 19.1 | 11_10_0_5 | 32629.5 | 6.3 |
| WT40 | 32637.9 | 60.5 | 12_10_1_4 | 32646.5 | 0.6 |
| WT41 | 32659.6 | 39.0 | 10_11_0_5 | 32670.5 | 2.9 |
| WT42 | 32725.4 | 55.2 | 10_12_1_4 | 32728.5 | 4.9 |
| WT43 | 32747.7 | 58.2 | 12_12_1_3 | 32761.5 | 5.8 |
| WT44 | 32768.8 | 4.1 | 11_10_1_5 | 32775.5 | 1.3 |
| WT45 | 32826.8 | 1.7 | 11_11_0_5 | 32832.5 | 2.3 |
| WT46 | 32851.5 | 10.7 | 13_11_0_4 | 32865.5 | 6.1 |
| WT47 | 32873.7 | 4.3 | 12_14_1_2 | 32876.6 | 5.1 |
| WT48 | 32927.5 | 49.5 | 12_10_1_5 | 32937.6 | 2.1 |
| WT49 | 32949.8 | 15.6 | 10_11_0_6 | 32961.6 | 3.8 |
| WT50 | 33293.5 | 34.6 | 13_11_1_5 | 33302.7 | 1.2 |
| WT51 | 33583.8 | 9.4 | 13_11_1_6 | 33593.8 | 2.0 |

**Table S.4.** Summary of molecular weights (MWs) of B.1.1.7 RBD glycoforms (Alpha, Ax) with respective relative abundances identified by ESI-MS and putative H_N_F_S combinations.

| RBD | Measured  MW (Da) | Relative abundance | H_N_F_S combination | Theoretical MW (Da) | Mass difference (Da) |
| --- | --- | --- | --- | --- | --- |
| A1 | 30894.6 | 6.4 | 9_10_0_0 | 30899.0 | 4.3 |
| A2 | 31080.7 | 17.1 | 9_8_2_1 | 31076.0 | 4.7 |
| A3 | 31137.8 | 5.3 | 9_9_3_0 | 31134.1 | 3.7 |
| A4 | 31160.2 | 3.2 | 9_7_2_2 | 31164.0 | 3.8 |
| A5 | 31184.9 | 3.7 | 9_10_0_1 | 31190.1 | 5.2 |
| A6 | 31243.1 | 13.0 | 10_8_2_1 | 31238.1 | 5.1 |
| A7 | 31265.9 | 10.9 | 10_11_0_0 | 31264.1 | 1.8 |
| A8 | 31300.5 | 4.5 | 10_9_3_0 | 31296.1 | 4.4 |
| A9 | 31319.6 | 3.4 | 8_10_2_1 | 31320.1 | 0.5 |
| A10 | 31346.6 | 2.6 | 10_10_0_1 | 31352.1 | 5.5 |
| A11 | 31369.5 | 19.4 | 11_10_1_0 | 31369.1 | 0.4 |
| A12 | 31405.1 | 26.4 | 8_9_2_2 | 31408.1 | 3.0 |
| A13 | 31427.3 | 13.6 | 11_11_0_0 | 31426.1 | 1.1 |
| A14 | 31449.4 | 4.9 | 7_10_4_1 | 31450.2 | 0.8 |
| A15 | 31482.0 | 4.6 | 9_10_2_1 | 31482.2 | 0.1 |
| A16 | 31505.7 | 3.4 | 7_11_3_1 | 31507.2 | 1.5 |
| A17 | 31529.1 | 30.9 | 12_10_1_0 | 31531.2 | 2.0 |
| A18 | 31553.5 | 35.3 | 10_11_0_1 | 31555.2 | 1.7 |
| A19 | 31590.8 | 27.8 | 12_11_0_0 | 31588.2 | 2.6 |
| A20 | 31611.6 | 16.5 | 8_10_2_2 | 31611.2 | 0.4 |
| A21 | 31658.9 | 9.8 | 11_10_3_0 | 31661.2 | 2.3 |
| A22 | 31693.9 | 8.8 | 13_10_1_0 | 31693.2 | 0.6 |
| A23 | 31715.2 | 96.9 | 11_11_0_1 | 31717.2 | 2.0 |
| A24 | 31737.6 | 13.7 | 12_11_1_0 | 31734.3 | 3.3 |
| A25 | 31773.3 | 19.8 | 9_10_2_2 | 31773.3 | 0.1 |
| A26 | 31793.9 | 13.4 | 12_12_0_0 | 31791.3 | 2.7 |
| A27 | 31819.9 | 31.4 | 12_10_1_1 | 31822.3 | 2.4 |
| A28 | 31844.4 | 20.4 | 10_11_0_2 | 31846.3 | 1.9 |
| A29 | 31880.1 | 13.3 | 12_11_0_1 | 31879.3 | 0.8 |
| A30 | 31901.1 | 78.6 | 8_10_4_2 | 31903.3 | 2.3 |
| A31 | 31920.8 | 4.8 | 11_12_0_1 | 31920.3 | 0.5 |
| A32 | 31937.3 | 2.7 | 12_12_1_0 | 31937.3 | 0.1 |
| A33 | 31957.6 | 10.8 | 13_12_0_0 | 31953.3 | 4.2 |
| A34 | 31978.1 | 10.0 | 9_11_2_2 | 31976.3 | 1.7 |
| A35 | 31006.1 | 100.0 | 11_11_0_2 | 31008.3 | 2.2 |
| A36 | 32027.7 | 5.2 | 12_11_3_0 | 32026.4 | 1.3 |
| A37 | 32061.8 | 4.9 | 9_10_2_3 | 32064.4 | 2.6 |
| A38 | 32081.4 | 53.5 | 12_12_0_1 | 32082.4 | 1.0 |
| A39 | 32109.6 | 10.5 | 12_10_1_2 | 32113.4 | 3.8 |
| A40 | 32137.5 | 13.8 | 10_11_2_2 | 32138.4 | 0.9 |
| A41 | 32191.8 | 82.8 | 13_11_3_0 | 32188.4 | 3.4 |
| A42 | 32212.7 | 3.2 | 11_12_0_2 | 32211.4 | 1.3 |
| A43 | 32227.5 | 2.6 | 12_12_1_1 | 32228.4 | 1.0 |
| A44 | 32266.5 | 39.1 | 9_11_2_3 | 32267.4 | 0.9 |
| A45 | 32297.9 | 48.2 | 11_11_0_3 | 32299.4 | 1.5 |
| A46 | 32372.0 | 53.5 | 12_12_0_2 | 32373.5 | 1.5 |
| A47 | 32393.9 | 5.0 | 11_10_4_2 | 32389.5 | 4.4 |
| A48 | 32446.4 | 31.2 | 11_11_1_3 | 32445.5 | 0.9 |
| A49 | 32483.1 | 35.2 | 13_11_3_1 | 32479.5 | 3.6 |
| A50 | 32504.3 | 4.7 | 11_12_2_2 | 32503.5 | 0.7 |
| A51 | 32557.2 | 43.9 | 14_12_3_0 | 32553.6 | 3.7 |
| A52 | 32590.0 | 10.0 | 11_11_2_3 | 32591.5 | 1.6 |
| A53 | 32631.8 | 25.2 | 10_12_2_3 | 32632.6 | 0.8 |
| A54 | 32663.5 | 34.6 | 12_12_0_3 | 32664.6 | 1.1 |
| A55 | 32707.1 | 4.5 | 11_13_2_2 | 32706.6 | 0.5 |
| A56 | 32737.2 | 44.7 | 11_11_1_4 | 32736.6 | 0.6 |
| A57 | 32774.4 | 6.2 | 13_11_3_2 | 32770.6 | 3.8 |
| A58 | 32811.6 | 21.6 | 12_12_1_3 | 32810.6 | 0.9 |
| A59 | 32848.2 | 24.6 | 14_12_3_1 | 32844.7 | 3.6 |
| A60 | 32869.8 | 4.8 | 12_13_2_2 | 32868.7 | 1.2 |
| A61 | 32887.8 | 3.5 | 13_13_1_2 | 32884.7 | 3.1 |
| A62 | 32922.7 | 34.7 | 10_12_2_4 | 32923.7 | 1.0 |
| A63 | 32954.7 | 11.5 | 12_12_0_4 | 32955.7 | 1.0 |
| A64 | 32998.0 | 16.2 | 11_13_2_3 | 32997.7 | 0.3 |
| A65 | 33028.4 | 28.5 | 11_11_1_5 | 33027.7 | 0.7 |
| A66 | 33050.4 | 3.3 | 12_11_4_3 | 33045.7 | 4.7 |
| A67 | 33072.5 | 4.0 | 12_14_2_2 | 33071.7 | 0.7 |
| A68 | 33102.6 | 30.2 | 12_12_1_4 | 33101.7 | 0.8 |
| A69 | 33138.5 | 8.1 | 14_12_3_2 | 33135.7 | 2.8 |
| A70 | 33176.8 | 16.5 | 13_13_1_3 | 33175.8 | 1.0 |
| A71 | 33213.2 | 22.7 | 15_13_3_1 | 33207.8 | 5.5 |
| A72 | 33250.3 | 3.8 | 12_12_2_4 | 33247.8 | 2.5 |
| A73 | 33288.0 | 23.1 | 16_14_3_0 | 33283.8 | 4.1 |
| A74 | 33320.0 | 12.6 | 11_11_3_5 | 33319.8 | 0.2 |
| A75 | 33362.3 | 13.4 | 12_14_2_3 | 33362.8 | 0.5 |
| A76 | 33393.8 | 21.4 | 12_12_1_5 | 33392.8 | 1.0 |
| A77 | 33467.9 | 24.3 | 13_13_1_4 | 33466.8 | 1.0 |
| A78 | 33504.4 | 8.7 | 15_13_3_2 | 33500.9 | 3.5 |
| A79 | 33542.5 | 11.6 | 14_14_1_3 | 33540.9 | 1.6 |
| A80 | 33578.2 | 15.7 | 16_14_3_1 | 33574.9 | 3.3 |
| A81 | 33613.2 | 4.6 | 13_13_0_5 | 33611.9 | 1.3 |
| A82 | 33653.0 | 18.2 | 12_14_2_4 | 33653.9 | 0.9 |
| A83 | 33685.1 | 8.2 | 14_14_2_3 | 33686.9 | 1.9 |
| A84 | 33727.9 | 11.9 | 11_13_3_5 | 33726.0 | 1.9 |
| A85 | 33759.2 | 17.4 | 13_13_1_5 | 33757.9 | 1.3 |
| A86 | 33833.4 | 16.8 | 14_14_1_4 | 33832.0 | 1.4 |
| A87 | 33868.9 | 7.3 | 16_14_1_3 | 33865.0 | 3.9 |
| A88 | 33907.0 | 11.8 | 13_13_0_6 | 33903.0 | 4.0 |
| A89 | 33944.0 | 12.4 | 13_13_0_6 | 33945.0 | 1.0 |
| A90 | 34018.6 | 14.8 | 12_14_2_5 | 34018.0 | 0.6 |
| A91 | 34051.1 | 10.3 | 13_13_3_5 | 34050.1 | 1.0 |
| A92 | 34125.0 | 11.1 | 14_14_1_5 | 34123.1 | 2.0 |
| A93 | 34198.1 | 9.5 | 13_13_0_7 | 34194.1 | 4.0 |
| A94 | 34309.0 | 7.9 | 13_15_0_6 | 34309.1 | 0.2 |
| A95 | 34599.3 | 3.5 | 13_15_2_6 | 34601.3 | 2.0 |
| A96 | 34823.3 | 2.2 | 16_13_1_7 | 34826.3 | 3.0 |
| A97 | 34886.3 | 8.7 | 16_14_2_6 | 34884.3 | 2.0 |

**Table S.5.** Summary of molecular weights (MWs) of B.1.351 RBD glycoforms (Beta, Bx) with respective relative abundances identified by ESI-MS and putative H_N_F_S combinations.

| RBD | Measured  MW (Da) | Relative abundance | H_N_F_S combination | Theoretical MW (Da) | Mass difference (Da) |
| --- | --- | --- | --- | --- | --- |
| B1 | 31247.7 | 10.6 | 9_9_0_2 | 31255.0 | 7.3 |
| B2 | 31352.5 | 18.2 | 10_8_1_2 | 31360.0 | 7.6 |
| B3 | 31376.3 | 12.9 | 9_11_0_1 | 31370.0 | 6.2 |
| B4 | 31408.6 | 25.9 | 11_11_0_0 | 31403.1 | 5.5 |
| B5 | 31433.0 | 16.9 | 11_9_1_1 | 31434.0 | 1.0 |
| B6 | 31512.7 | 19.6 | 9_11_1_1 | 31516.1 | 3.4 |
| B7 | 31538.1 | 62.3 | 9_9_0_3 | 31546.1 | 8.0 |
| B8 | 31559.8 | 7.0 | 10_9_1_2 | 31563.1 | 3.2 |
| B9 | 31595.6 | 34.7 | 9_10_1_2 | 31604.1 | 8.6 |
| B10 | 31616.5 | 9.9 | 10_10_0_2 | 31620.1 | 3.6 |
| B11 | 31641.7 | 11.0 | 11_10_1_1 | 31637.1 | 4.6 |
| B12 | 31669.2 | 15.4 | 13_10_1_0 | 31670.1 | 0.9 |
| B13 | 31700.2 | 100.0 | 10_9_0_3 | 31708.1 | 7.9 |
| B14 | 31723.8 | 64.4 | 11_9_1_2 | 31725.1 | 1.3 |
| B15 | 31758.7 | 10.9 | 10_10_1_2 | 31766.2 | 7.5 |
| B16 | 31778.5 | 16.2 | 11_10_0_2 | 31782.2 | 3.7 |
| B17 | 31803.0 | 12.5 | 12_10_1_1 | 31799.2 | 3.8 |
| B18 | 31829.5 | 42.9 | 10_11_2_1 | 31824.2 | 5.3 |
| B19 | 31886.1 | 84.9 | 12_9_1_2 | 31887.2 | 1.1 |
| B20 | 31906.0 | 32.1 | 10_10_0_3 | 31911.2 | 5.2 |
| B21 | 31962.0 | 22.6 | 10_11_1_2 | 31969.3 | 7.2 |
| B22 | 31991.3 | 58.7 | 10_9_2_3 | 31000.2 | 8.9 |
| B23 | 32014.8 | 31.8 | 11_9_1_3 | 32016.2 | 1.5 |
| B24 | 32066.4 | 53.9 | 11_10_0_3 | 32073.3 | 6.9 |
| B25 | 32089.4 | 36.3 | 12_10_1_2 | 32090.3 | 0.9 |
| B26 | 32142.2 | 10.0 | 12_11_0_2 | 32147.3 | 5.1 |
| B27 | 32177.0 | 45.9 | 12_9_1_3 | 32178.3 | 1.3 |
| B28 | 32196.7 | 21.8 | 10_10_0_4 | 32202.3 | 5.6 |
| B29 | 32251.5 | 48.1 | 13_10_1_2 | 32252.3 | 0.9 |
| B30 | 32271.1 | 10.1 | 11_11_0_3 | 32276.3 | 5.3 |
| B31 | 32326.4 | 14.9 | 14_11_1_1 | 32326.4 | 0.1 |
| B32 | 32356.4 | 33.3 | 11_10_0_4 | 32364.4 | 8.0 |
| B33 | 32379.5 | 27.6 | 12_10_1_3 | 32381.4 | 1.9 |
| B34 | 32431.3 | 48.8 | 12_11_0_3 | 32438.4 | 7.1 |
| B35 | 32454.7 | 29.5 | 11_9_2_4 | 32453.4 | 1.3 |
| B36 | 32541.6 | 24.4 | 13_10_1_3 | 32543.4 | 1.8 |
| B37 | 32560.9 | 20.8 | 11_11_0_4 | 32567.4 | 6.6 |
| B38 | 32617.1 | 43.1 | 14_11_1_2 | 32617.5 | 0.4 |
| B39 | 32637.1 | 9.6 | 12_12_0_3 | 32641.5 | 4.4 |
| B40 | 32722.1 | 35.2 | 12_11_0_4 | 32729.5 | 7.4 |
| B41 | 32744.8 | 17.2 | 13_11_1_3 | 32746.5 | 1.7 |
| B42 | 32796.8 | 34.1 | 11_10_1_5 | 32801.5 | 4.7 |
| B43 | 32820.3 | 13.6 | 12_10_2_4 | 32818.5 | 1.8 |
| B44 | 32927.0 | 9.0 | 12_12_0_4 | 32932.6 | 5.6 |
| B45 | 32983.0 | 30.0 | 13_10_2_4 | 32980.6 | 2.4 |
| B46 | 33109.9 | 10.9 | 12_10_2_5 | 33109.6 | 0.3 |
| B47 | 33162.3 | 26.9 | 12_11_1_5 | 33166.6 | 4.4 |
| B48 | 33273.5 | 21.2 | 13_10_2_5 | 33271.7 | 1.8 |
| B49 | 33348.1 | 27.5 | 14_11_2_4 | 33345.7 | 2.4 |
| B50 | 33452.7 | 24.3 | 12_11_1_6 | 33457.7 | 5.1 |
| B51 | 33528.0 | 25.5 | 13_12_1_5 | 33531.8 | 3.8 |
| B52 | 33637.9 | 17.3 | 14_11_2_5 | 33636.8 | 1.1 |
| B53 | 33713.1 | 20.7 | 13_10_3_6 | 33708.8 | 4.3 |
| B54 | 33817.9 | 21.0 | 13_12_1_6 | 33822.9 | 5.0 |
| B55 | 33893.0 | 17.0 | 14_13_1_5 | 33896.9 | 3.9 |
| B56 | 34003.3 | 13.8 | 13_10_3_7 | 33999.9 | 3.4 |
| B57 | 34183.0 | 10.0 | 14_13_1_6 | 34188.0 | 5.0 |
| B58 | 34548.7 | 7.7 | 15_14_1_6 | 34553.1 | 4.5 |
| B59 | 34693.1 | 13.5 | 15_14_2_6 | 34699.2 | 6.1 |

**Table S.6.** Summary of molecular weights (MWs) of B.1.351.2 RBD glycoforms (Delta, Dx) with respective relative abundances identified by ESI-MS and putative H_N_F_S combinations.

| RBD | Measured  MW (Da) | Relative abundance | H_N_F_S combination | Theoretical MW (Da) | Mass difference (Da) |
| --- | --- | --- | --- | --- | --- |
| D1 | 31073.5 | 20.9 | 8_11_1_0 | 31079.0 | 5.5 |
| D2 | 31094.3 | 1.0 | 9_11_0_0 | 31095.0 | 0.7 |
| D3 | 31259.9 | 24.3 | 10_11_0_0 | 31257.1 | 2.8 |
| D4 | 31363.7 | 28.6 | 8_11_1_1 | 31370.1 | 6.4 |
| D5 | 31423.0 | 23.4 | 11_11_0_0 | 31419.1 | 3.9 |
| D6 | 31526.2 | 26.6 | 9_11_1_1 | 31532.2 | 5.9 |
| D7 | 31548.3 | 25.4 | 10_11_0_1 | 31548.2 | 0.2 |
| D8 | 31654.2 | 29.4 | 8_11_1_2 | 31661.2 | 7.0 |
| D9 | 31690.5 | 23.9 | 10_11_1_1 | 31694.2 | 3.7 |
| D10 | 31711.4 | 39.3 | 11_11_0_1 | 31710.2 | 1.2 |
| D11 | 31733.5 | 16.1 | 9_12_1_1 | 31735.2 | 1.7 |
| D12 | 31813.8 | 33.6 | 10_13_1_0 | 31809.3 | 4.5 |
| D13 | 31838.0 | 31.0 | 10_11_0_2 | 31839.3 | 1.3 |
| D14 | 31875.9 | 13.2 | 12_11_2_0 | 31873.3 | 2.6 |
| D15 | 31895.2 | 25.7 | 10_12_1_1 | 31897.3 | 2.1 |
| D16 | 31979.4 | 30.5 | 10_11_1_2 | 31985.3 | 5.9 |
| D17 | 31000.6 | 60.7 | 11_11_0_2 | 31001.3 | 0.8 |
| D18 | 32022.0 | 26.0 | 9_12_1_2 | 32026.3 | 4.3 |
| D19 | 32079.6 | 25.4 | 12_12_0_1 | 32075.3 | 4.2 |
| D20 | 32104.2 | 49.0 | 10_13_1_1 | 32100.4 | 3.8 |
| D21 | 32127.5 | 29.3 | 10_11_0_3 | 32130.3 | 2.8 |
| D22 | 32164.9 | 4.6 | 12_11_0_2 | 32163.4 | 1.6 |
| D23 | 32185.4 | 44.8 | 10_12_1_2 | 32188.4 | 3.0 |
| D24 | 32206.8 | 5.0 | 11_12_0_2 | 32204.4 | 2.4 |
| D25 | 32290.9 | 90.6 | 11_11_0_3 | 32292.4 | 1.5 |
| D26 | 32312.4 | 32.4 | 9_12_1_3 | 32317.4 | 5.1 |
| D27 | 32348.4 | 26.0 | 11_12_1_2 | 32350.4 | 2.1 |
| D28 | 32369.1 | 34.1 | 12_12_0_2 | 32366.4 | 2.7 |
| D29 | 32394.4 | 45.5 | 10_13_1_2 | 32391.5 | 3.0 |
| D30 | 32475.9 | 50.0 | 10_12_1_3 | 32479.5 | 3.5 |
| D31 | 32496.5 | 22.2 | 11_12_0_3 | 32495.5 | 1.0 |
| D32 | 32553.6 | 0.7 | 11_13_1_2 | 32553.5 | 0.1 |
| D33 | 32581.7 | 100.0 | 11_11_0_4 | 32583.5 | 1.8 |
| D34 | 32603.7 | 34.7 | 12_11_3_2 | 32601.5 | 2.2 |
| D35 | 32638.2 | 0.3 | 11_12_1_3 | 32641.5 | 3.3 |
| D36 | 32658.4 | 41.3 | 12_12_0_3 | 32657.5 | 0.8 |
| D37 | 32683.3 | 21.8 | 10_13_1_3 | 32682.6 | 0.7 |
| D38 | 32766.4 | 41.9 | 10_12_1_4 | 32770.6 | 4.2 |
| D39 | 32843.0 | 8.0 | 11_13_1_3 | 32844.6 | 1.7 |
| D40 | 32873.4 | 68.5 | 11_11_0_5 | 32874.6 | 1.2 |
| D41 | 32895.2 | 18.5 | 12_11_3_3 | 32892.6 | 2.6 |
| D42 | 32948.2 | 45.1 | 12_12_0_4 | 32948.6 | 0.4 |
| D43 | 33238.9 | 42.5 | 12_12_0_5 | 33239.7 | 0.8 |
| D44 | 33313.9 | 30.2 | 13_13_0_4 | 33313.8 | 0.1 |
| D45 | 33604.6 | 37.1 | 13_13_0_5 | 33604.9 | 0.2 |
| D46 | 33896.1 | 29.5 | 13_13_0_6 | 33896.0 | 0.2 |
| D47 | 33969.9 | 9.2 | 12_12_3_6 | 33969.0 | 0.9 |
| D48 | 34186.4 | 21.6 | 13_13_0_7 | 34187.0 | 0.7 |
| D49 | 34260.3 | 9.0 | 14_14_0_6 | 34261.1 | 0.7 |

**Table S.7.** Summary of molecular weights (MWs) of B.1.1.529 RBD glycoforms (Omicron, Ox) with respective relative abundances identified by ESI-MS and putative H_N_F_S combinations.

| RBD | Measured  MW (Da) | Relative abundance | H_N_F_S combination | Theoretical MW (Da) | Mass difference (Da) |
| --- | --- | --- | --- | --- | --- |
| O1 | 32489.2 | 40.2 | 10_11_1_3 | 32487.6 | 1.5 |
| O2 | 32538.4 | 45.7 | 11_9_3_3 | 32535.7 | 2.7 |
| O3 | 32559.4 | 5.6 | 11_12_1_2 | 32561.7 | 2.3 |
| O4 | 32576.0 | 23.0 | 10_10_1_4 | 32575.7 | 0.3 |
| O5 | 32595.1 | 74.6 | 11_10_2_3 | 32592.7 | 2.4 |
| O6 | 32615.4 | 31.4 | 12_10_1_3 | 32608.7 | 6.7 |
| O7 | 32651.1 | 29.0 | 11_11_1_3 | 32649.7 | 1.4 |
| O8 | 32670.2 | 24.9 | 12_11_2_2 | 32666.7 | 3.4 |
| O9 | 32702.3 | 42.5 | 11_12_0_3 | 32706.7 | 4.4 |
| O10 | 32723.8 | 87.1 | 10_10_2_4 | 32721.7 | 2.1 |
| O11 | 32744.0 | 2.5 | 11_10_3_3 | 32738.7 | 5.3 |
| O12 | 32761.0 | 11.3 | 12_10_2_3 | 32754.7 | 6.3 |
| O13 | 32779.8 | 53.6 | 10_11_1_4 | 32778.7 | 1.1 |
| O14 | 32798.7 | 23.0 | 11_11_2_3 | 32795.8 | 2.9 |
| O15 | 32829.1 | 62.1 | 11_9_3_4 | 32826.8 | 2.3 |
| O16 | 32852.5 | 27.6 | 11_12_1_3 | 32852.8 | 0.3 |
| O17 | 32885.7 | 85.7 | 11_10_2_4 | 32883.8 | 2.0 |
| O18 | 32905.7 | 29.7 | 12_10_3_3 | 32900.8 | 4.9 |
| O19 | 32942.2 | 12.1 | 11_11_1_4 | 32940.8 | 1.4 |
| O20 | 32960.8 | 40.8 | 12_11_2_3 | 32957.8 | 3.0 |
| O21 | 33014.9 | 100.0 | 12_12_1_3 | 33014.8 | 0.1 |
| O22 | 33035.6 | 25.5 | 11_10_3_4 | 33029.8 | 5.8 |
| O23 | 33070.2 | 52.1 | 12_13_0_3 | 33071.9 | 1.6 |
| O24 | 33089.8 | 52.7 | 11_11_2_4 | 33086.9 | 2.9 |
| O25 | 33120.4 | 52.4 | 13_11_2_3 | 33119.9 | 0.6 |
| O26 | 33144.7 | 28.3 | 11_12_1_4 | 33143.9 | 0.8 |
| O27 | 33177.5 | 50.7 | 11_10_2_5 | 33174.9 | 2.6 |
| O28 | 33195.9 | 33.2 | 12_10_3_4 | 33191.9 | 4.0 |
| O29 | 33217.4 | 13.2 | 12_13_1_3 | 33217.9 | 0.5 |
| O30 | 33233.9 | 1.6 | 11_11_1_5 | 33231.9 | 2.0 |
| O31 | 33251.6 | 58.4 | 12_11_2_4 | 33248.9 | 2.7 |
| O32 | 33271.2 | 15.4 | 13_11_3_3 | 33265.9 | 5.2 |
| O33 | 33306.1 | 83.2 | 12_12_1_4 | 33305.9 | 0.2 |
| O34 | 33326.7 | 28.9 | 13_12_2_3 | 33322.9 | 3.7 |
| O35 | 33361.5 | 30.3 | 12_13_0_4 | 33362.9 | 1.4 |
| O36 | 33380.8 | 61.4 | 13_13_1_3 | 33380.0 | 0.9 |
| O37 | 33402.5 | 0.3 | 14_13_0_3 | 33396.0 | 6.5 |
| O38 | 33435.9 | 27.5 | 11_12_1_5 | 33435.0 | 0.9 |
| O39 | 33454.8 | 15.1 | 12_12_2_4 | 33452.0 | 2.8 |
| O40 | 33486.2 | 24.5 | 14_12_0_4 | 33484.0 | 2.2 |
| O41 | 33508.3 | 18.3 | 12_13_1_4 | 33509.0 | 0.7 |
| O42 | 33542.1 | 37.0 | 12_11_2_5 | 33540.0 | 2.1 |
| O43 | 33561.5 | 24.8 | 15_13_0_3 | 33558.0 | 3.5 |
| O44 | 33597.5 | 25.2 | 12_12_1_5 | 33597.0 | 0.5 |
| O45 | 33617.3 | 33.1 | 13_12_2_4 | 33614.0 | 3.2 |
| O46 | 33671.8 | 47.2 | 13_13_1_4 | 33671.1 | 0.7 |
| O47 | 33691.7 | 16.4 | 14_13_0_4 | 33687.1 | 4.6 |
| O48 | 33726.3 | 13.7 | 13_14_0_4 | 33728.1 | 1.7 |
| O49 | 33745.7 | 35.3 | 12_12_2_5 | 33743.1 | 2.6 |
| O50 | 33776.2 | 16.1 | 14_12_2_4 | 33776.1 | 0.1 |
| O51 | 33800.4 | 17.0 | 12_13_1_5 | 33800.1 | 0.3 |
| O52 | 33833.8 | 3.0 | 14_13_1_4 | 33833.1 | 0.7 |
| O53 | 33851.7 | 28.4 | 15_13_0_4 | 33849.1 | 2.6 |
| O54 | 33874.5 | 10.3 | 13_14_1_4 | 33874.1 | 0.4 |
| O55 | 33907.8 | 21.7 | 13_12_2_5 | 33905.1 | 2.7 |
| O56 | 33927.2 | 14.4 | 14_12_3_4 | 33922.1 | 5.0 |
| O57 | 33963.1 | 27.1 | 13_13_1_5 | 33962.2 | 0.9 |
| O58 | 33982.3 | 13.7 | 14_13_0_5 | 33978.1 | 4.2 |
| O59 | 34037.0 | 32.6 | 14_14_1_4 | 34036.2 | 0.8 |
| O60 | 34058.2 | 3.7 | 13_12_3_5 | 34051.2 | 7.0 |
| O61 | 34090.9 | 8.8 | 12_13_1_6 | 34091.2 | 0.3 |
| O62 | 34110.9 | 11.6 | 13_13_2_5 | 34108.2 | 2.7 |
| O63 | 34142.7 | 22.5 | 15_13_0_5 | 34140.2 | 2.5 |
| O64 | 34165.6 | 5.1 | 13_14_1_5 | 34165.2 | 0.4 |
| O65 | 34199.3 | 6.6 | 15_14_1_4 | 34198.2 | 1.1 |
| O66 | 34217.1 | 20.9 | 14_12_1_6 | 34212.2 | 4.8 |
| O67 | 34274.9 | 20.8 | 14_13_2_5 | 34270.3 | 4.6 |
| O68 | 34327.5 | 26.7 | 14_14_1_5 | 34327.3 | 0.2 |
| O69 | 34346.6 | 15.5 | 13_12_3_6 | 34342.3 | 4.3 |
| O70 | 34384.5 | 0.9 | 13_10_4_7 | 34373.3 | 11.2 |
| O71 | 34402.4 | 23.3 | 13_13_2_6 | 34399.3 | 3.1 |
| O72 | 34457.1 | 9.0 | 13_14_1_6 | 34456.3 | 0.8 |
| O73 | 34506.7 | 18.8 | 14_12_1_7 | 34503.3 | 3.4 |
| O74 | 34529.7 | 3.1 | 12_13_4_6 | 34529.4 | 0.3 |
| O75 | 34564.3 | 25.0 | 14_13_0_7 | 34560.3 | 4.0 |
| O76 | 34620.0 | 26.6 | 14_14_1_6 | 34618.4 | 1.6 |
| O77 | 34638.4 | 23.0 | 15_14_2_5 | 34635.4 | 3.0 |
| O78 | 34658.8 | 12.7 | 16_14_3_4 | 34652.4 | 6.4 |
| O79 | 34768.4 | 0.9 | 14_14_2_6 | 34764.4 | 4.0 |
| O80 | 34798.6 | 29.8 | 14_12_1_8 | 34794.4 | 4.2 |
| O81 | 34840.4 | 0.5 | 13_13_3_7 | 34836.5 | 4.0 |
| O82 | 34858.9 | 20.4 | 14_13_0_8 | 34851.4 | 7.5 |
| O83 | 35019.8 | 14.5 | 15_13_0_8 | 35013.5 | 6.3 |
| O84 | 35041.2 | 10.5 | 13_14_3_7 | 35039.5 | 1.7 |
| O85 | 35220.7 | 23.8 | 15_14_0_8 | 35216.6 | 4.1 |
| O86 | 35312.3 | 2.6 | 15_13_2_8 | 35305.6 | 6.7 |
| O87 | 35354.0 | 19.8 | 15_14_1_8 | 35362.6 | 8.6 |

**Table S.8.** Summary of molecular weights (MWs) of endoF3-treated WHu-Hu-1 RBD glycoforms (eWTx) with respective relative abundances identified by ESI-MS and putative H_N_F_S combinations.

| RBD | Measured  MW (Da) | Relative abundance | H_N_F_S combination | Theoretical MW (Da) | Mass difference (Da) | Number of trimmed N-glycans |
| --- | --- | --- | --- | --- | --- | --- |
| eWT1 | 28116.9 | 4.5 | 1_3_0_0 | 28123.9 | 7.0 | 2 |
| eWT2 | 28243.9 | 26.4 | 3_2_0_0 | 28244.9 | 1.0 | 2 |
| eWT3 | 28357.3 | 14.5 | 1_2_1_1 | 28358.0 | 0.6 | 2 |
| eWT4 | 28618.0 | 23.8 | 1_4_2_0 | 28619.1 | 1.0 | 2 |
| eWT5 | 28881.1 | 2.4 | 3_3_1_1 | 28885.1 | 4.0 | 2 |
| eWT6 | 29029.2 | 18.7 | 3_3_2_1 | 29031.2 | 2.1 | 2 |
| eWT7 | 29143.8 | 1.1 | 6_4_0_0 | 29137.2 | 6.5 | 1 |
| eWT8 | 29318.2 | 1.1 | 5_5_1_0 | 29324.3 | 6.1 | 1 |
| eWT9 | 29406.5 | 8.2 | 5_4_1_1 | 29412.3 | 5.8 | 1 |
| eWT10 | 29454.5 | 14.1 | 4_5_1_1 | 29453.4 | 1.2 | 1 |
| eWT11 | 29649.4 | 0.6 | 7_5_1_0 | 29648.4 | 1.0 | 1 |
| eWT12 | 29736.3 | 9.6 | 5_7_1_0 | 29730.5 | 5.8 | 1 |
| eWT13 | 29753.5 | 1.3 | 5_5_0_2 | 29760.5 | 6.9 | 1 |
| eWT14 | 30084.9 | 1.3 | 7_5_2_1 | 30085.6 | 0.7 | 1 |
| eWT15 | 30103.4 | 10.3 | 5_6_1_2 | 30109.6 | 6.2 | 1 |
| eWT16 | 31033.0 | 27.6 | 8_6_2_3 | 31032.9 | 0.1 | 1 |
| eWT17 | 31216.4 | 17.2 | 7_7_1_4 | 31219.0 | 2.5 | 1 |
| eWT18 | 31416.8 | 8.0 | 7_8_1_4 | 31422.0 | 5.2 | 1 |
| eWT19 | 31437.4 | 0.9 | 8_8_0_4 | 31438.0 | 0.7 | 1 |
| eWT20 | 31482.3 | 9.0 | 7_9_2_3 | 31480.1 | 2.2 | 1 |
| eWT21 | 31505.7 | 38.4 | 9_9_0_3 | 31512.1 | 6.4 | 0 |
| eWT22 | 31543.3 | 41.0 | 9_7_1_4 | 31543.1 | 0.2 | 1 |
| eWT23 | 31564.8 | 55.2 | 7_8_2_4 | 31568.1 | 3.3 | 1 |
| eWT24 | 31607.4 | 2.2 | 11_10_3_0 | 31604.1 | 3.2 | 0 |
| eWT25 | 31644.1 | 27.3 | 8_9_2_3 | 31642.1 | 2.0 | 1 |
| eWT26 | 31668.2 | 82.8 | 10_9_0_3 | 31674.1 | 5.9 | 0 |
| eWT27 | 31691.0 | 33.8 | 11_9_1_2 | 31691.1 | 0.2 | 0 |
| eWT28 | 31708.4 | 9.9 | 9_10_0_3 | 31715.2 | 6.7 | 0 |
| eWT29 | 31727.6 | 58.6 | 10_10_1_2 | 31732.2 | 4.5 | 0 |
| eWT30 | 31748.3 | 42.7 | 11_10_0_2 | 31748.2 | 0.1 | 0 |
| eWT31 | 31770.9 | 29.2 | 12_10_3_0 | 31766.2 | 4.7 | 0 |
| eWT32 | 31795.9 | 25.7 | 10_11_0_2 | 31789.2 | 6.7 | 0 |
| eWT33 | 31833.6 | 35.0 | 11_9_2_2 | 31837.2 | 3.6 | 0 |
| eWT34 | 31854.4 | 95.9 | 9_10_1_3 | 31861.2 | 6.8 | 0 |
| eWT35 | 31872.6 | 14.1 | 10_10_2_2 | 31878.2 | 5.6 | 0 |
| eWT36 | 31893.5 | 1.5 | 11_10_1_2 | 31894.2 | 0.7 | 0 |
| eWT37 | 31912.9 | 51.6 | 12_10_0_2 | 31910.2 | 2.6 | 0 |
| eWT38 | 31933.2 | 26.1 | 10_11_1_2 | 31935.3 | 2.0 | 0 |
| eWT39 | 31958.1 | 100.0 | 11_11_0_2 | 31951.2 | 6.9 | 0 |
| eWT40 | 31980.4 | 28.5 | 13_11_0_1 | 31984.3 | 3.9 | 0 |
| eWT41 | 32017.7 | 36.8 | 10_10_1_3 | 32023.3 | 5.5 | 0 |
| eWT42 | 32060.9 | 6.2 | 12_10_1_2 | 32056.3 | 4.6 | 0 |

**Table S.9.** Summary of molecular weights (MWs) of endoF3-treated B.1.351.2 RBD glycoforms (eDx) with respective relative abundances identified by ESI-MS and putative H_N_F_S combinations.

| RBD | Measured  MW (Da) | Relative abundance | H_N_F_S combination | Theoretical MW (Da) | Mass difference (Da) | Number of trimmed N-glycans |
| --- | --- | --- | --- | --- | --- | --- |
| eD1 | 28681.8 | 2.2 | 4_3_0_0 | 28680.1 | 1.7 | 2 |
| eD2 | 28720.3 | 24.9 | 3_4_0_0 | 28721.1 | 0.8 | 2 |
| eD3 | 28741.8 | 35.3 | 2_2_2_1 | 28736.1 | 5.6 | 2 |
| eD4 | 28762.6 | 9.4 | 2_5_0_0 | 28762.2 | 0.5 | 2 |
| eD5 | 28782.3 | 33.5 | 1_3_2_1 | 28777.2 | 5.2 | 2 |
| eD6 | 28803.6 | 25.9 | 3_3_2_0 | 28810.2 | 6.5 | 2 |
| eD7 | 28826.2 | 19.8 | 4_3_1_0 | 28826.2 | 0.1 | 2 |
| eD8 | 28867.3 | 82.4 | 3_4_1_0 | 28867.2 | 0.1 | 2 |
| eD9 | 28888.2 | 52.6 | 4_4_0_0 | 28883.2 | 5.0 | 2 |
| eD10 | 28907.6 | 20.6 | 3_2_2_1 | 28898.2 | 9.4 | 2 |
| eD11 | 28927.8 | 100.0 | 3_5_0_0 | 28924.2 | 3.6 | 2 |
| eD12 | 28949.2 | 57.5 | 3_3_1_1 | 28955.2 | 6.0 | 2 |
| eD13 | 28970.6 | 38.3 | 4_3_0_1 | 28971.2 | 0.6 | 2 |
| eD14 | 28990.7 | 30.3 | 2_4_1_1 | 28996.2 | 5.5 | 2 |
| eD15 | 29011.6 | 32.3 | 3_4_0_1 | 29012.2 | 0.6 | 2 |
| eD16 | 29032.7 | 24.1 | 4_4_1_0 | 29029.2 | 3.5 | 2 |
| eD17 | 29053.6 | 52.1 | 3_2_1_2 | 29043.2 | 10.4 | 2 |
| eD18 | 29074.7 | 51.9 | 3_5_1_0 | 29070.3 | 4.4 | 2 |
| eD19 | 29095.1 | 15.7 | 3_3_2_1 | 29101.3 | 6.2 | 2 |
| eD20 | 29114.2 | 37.7 | 4_3_1_1 | 29117.3 | 3.1 | 2 |
| eD21 | 29135.1 | 26.0 | 2_4_0_2 | 29141.3 | 6.2 | 2 |
| eD22 | 29158.3 | 34.9 | 3_4_1_1 | 29158.3 | 0.0 | 2 |
| eD23 | 29179.7 | 25.3 | 4_4_2_0 | 29175.3 | 4.4 | 2 |
| eD24 | 29198.8 | 17.6 | 3_2_2_2 | 29189.3 | 9.6 | 2 |
| eD25 | 29219.2 | 43.7 | 3_5_0_1 | 29215.3 | 3.9 | 2 |
| eD26 | 29240.2 | 24.3 | 3_3_1_2 | 29246.3 | 6.1 | 2 |
| eD27 | 29261.4 | 16.4 | 4_3_2_1 | 29263.3 | 1.9 | 2 |
| eD28 | 29281.5 | 6.4 | 2_4_1_2 | 29287.3 | 5.9 | 2 |
| eD29 | 29301.3 | 1.9 | 3_4_2_1 | 29304.3 | 3.0 | 2 |
| eD30 | 29302.0 | 1.2 | 3_4_0_2 | 29303.3 | 1.3 | 2 |
| eD31 | 29344.9 | 24.8 | 5_4_2_0 | 29337.4 | 7.6 | 2 |
| eD32 | 29366.8 | 5.0 | 3_5_1_1 | 29361.4 | 5.5 | 2 |
| eD33 | 29405.7 | 10.1 | 4_3_1_2 | 29408.4 | 2.7 | 2 |
| eD34 | 29427.1 | 3.8 | 2_4_2_2 | 29433.4 | 6.3 | 2 |

**Table S.10.** Glycopeptide analysis of WT RBD

| N-site | Peptide sequence | N-glycan | Retention time (min) | Measured m/z | Charge state | Mass accuracy (ppm) | MS area |
| --- | --- | --- | --- | --- | --- | --- | --- |
| N331 | PNITNLCPFGEV | A2S2 | 18.0 | 1170.471 | 3 | -3.66 | 88472 |
| N331 | PNITNLCPFGEV | A2S2 | 18.0 | 1755.705 | 2 | -2.48 | 70771 |
| N331 | PNITNLCPFGEV | A1G1F | 18.2 | 1354.577 | 2 | 0.45 | 157724 |
| N331 | PNITNLCPFGEV | A1G0F | 18.3 | 1273.551 | 2 | 0.14 | 85745 |
| N331 | PNITNLCPFGEV | A1S1F | 18.6 | 1500.124 | 2 | 0.04 | 65793 |
| N331 | FPNIT | A2G2B | 27.8 | 1209.489 | 2 | -2.73 | 57370 |
| N343 | GEVFNATR | A2S1G1FB | 31.0 | 1579.132 | 2 | -3.43 | 44341 |
| N343 | NATRF | A2G2F | 6.3 | 1189.484 | 2 | 0.33 | 107145 |
| N343 | NATRF | A2G1F | 6.5 | 1107.955 | 2 | 0.30 | 46566 |
| N343 | NATRF | A2S1G1F | 7.0 | 1335.032 | 2 | 0.75 | 38217 |
| N343 | PFGEVFNATR | A2G2 | 13.5 | 1381.076 | 2 | -3.29 | 272764 |
| N343 | PFGEVFNATR | A2G1 | 13.6 | 1300.049 | 2 | -3.69 | 150239 |
| N343 | PFGEVFNATR | A2S1G1 | 13.7 | 1526.623 | 2 | -2.90 | 305875 |
| N343 | PFGEVFNATR | A2S1G0 | 13.8 | 1445.597 | 2 | -3.53 | 164686 |

**Table S.11.** Glycopeptide analysis of endoF3-treated WT RBD

| N-site | Peptide sequence | N-glycan | Retention time (min) | Measured m/z | Charge state | Mass accuracy (ppm) | MS area |
| --- | --- | --- | --- | --- | --- | --- | --- |
| N331 | PNITNLCPFGEVF | A1G0F | 18.9 | 1347.585 | 2 | -3.88 | 21591 |
| N331 | PNITNLCPFGE | A2G0FB | 19.5 | 1427.087 | 2 | -5.81 | 83634 |
| N331 | PNITNLCPFGEV | A1G1F | 19.6 | 1354.575 | 2 | -1.09 | 112872 |
| N331 | PNITNLCPFGEV | A1G0F | 19.7 | 1273.551 | 2 | -1.21 | 52498 |
| N331 | PNITNLCPFGEV | A1S1F | 19.9 | 1500.122 | 2 | -1.51 | 59378 |
| N331 | PNITNLCPF | A2G1 | 26.2 | 827.350 | 3 | 1.33 | 49111 |
| N331 | PNITNLCPF | A2G1 | 26.9 | 1240.524 | 2 | 2.81 | 25854 |
| N331 | PNITNL | A1G1 | 28.6 | 964.915 | 2 | 0.19 | 19156 |
| N343 | NATRF | A2G2F | 6.3 | 1189.483 | 2 | 0.54 | 32460 |
| N343 | NATRF | A2G1F | 6.4 | 1107.955 | 2 | 0.19 | 47037 |
| N343 | NATRF | A2G0F | 6.5 | 1027.430 | 2 | 0.50 | 23831 |
| N343 | NATRF | A2S1G1F | 7.5 | 1335.032 | 2 | 0.75 | 34621 |
| N343 | NATRF | A2S2F | 7.8 | 1480.580 | 2 | 1.09 | 24974 |
| N343 | NATRF | GnF | 8.2 | 957.453 | 1 | 0.28 | 17273 |
| N343 | NATRF | GnF | 8.2 | 479.230 | 2 | 0.53 | 255747 |
| N343 | NATR | A2G0 | 18.0 | 880.363 | 2 | -2.3 | 34690 |
| N343 | NATRFASVY | A2S1G1 | 9.4 | 736.560 | 4 | 4.72 | 103643 |
| N343 | GEVFNATR | GnF | 11.3 | 621.797 | 2 | 0.83 | 14586 |
| N343 | GEVFNATR | GnF | 15.5 | 414.868 | 3 | 3.39 | 15706 |
| N343 | GEVFNATR | GnF | 15.7 | 414.868 | 3 | 3.78 | 18198 |
| N343 | GEVFNATRF | GnF | 17.6 | 695.331 | 2 | 1.23 | 17392 |
| N343 | GEVFNATRF | A2G0F | 23.2 | 1243.537 | 2 | 4.76 | 17779 |

**Additional Figures**

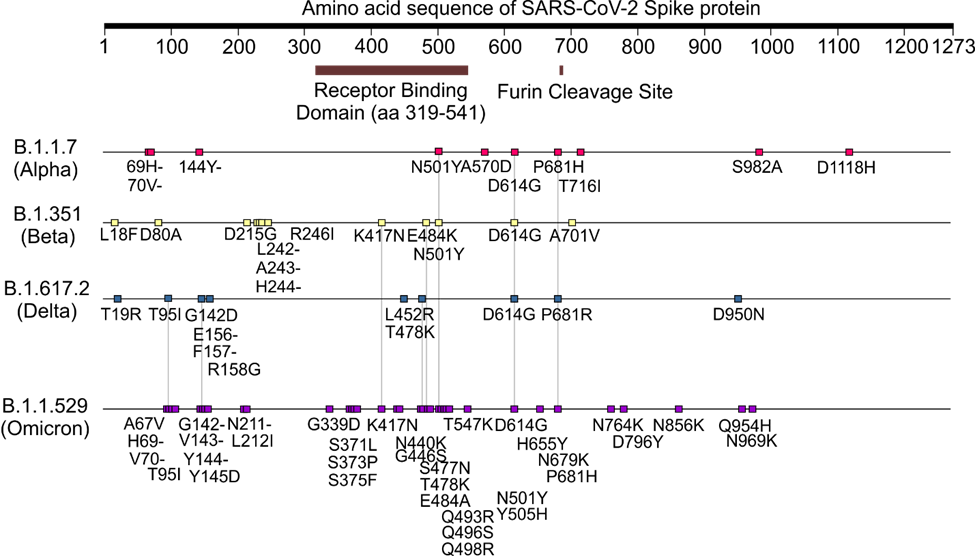

**Figure S.1** Amino acid sequence of SARS-CoV-2 S protein variants and mutations on RBD used in this study.

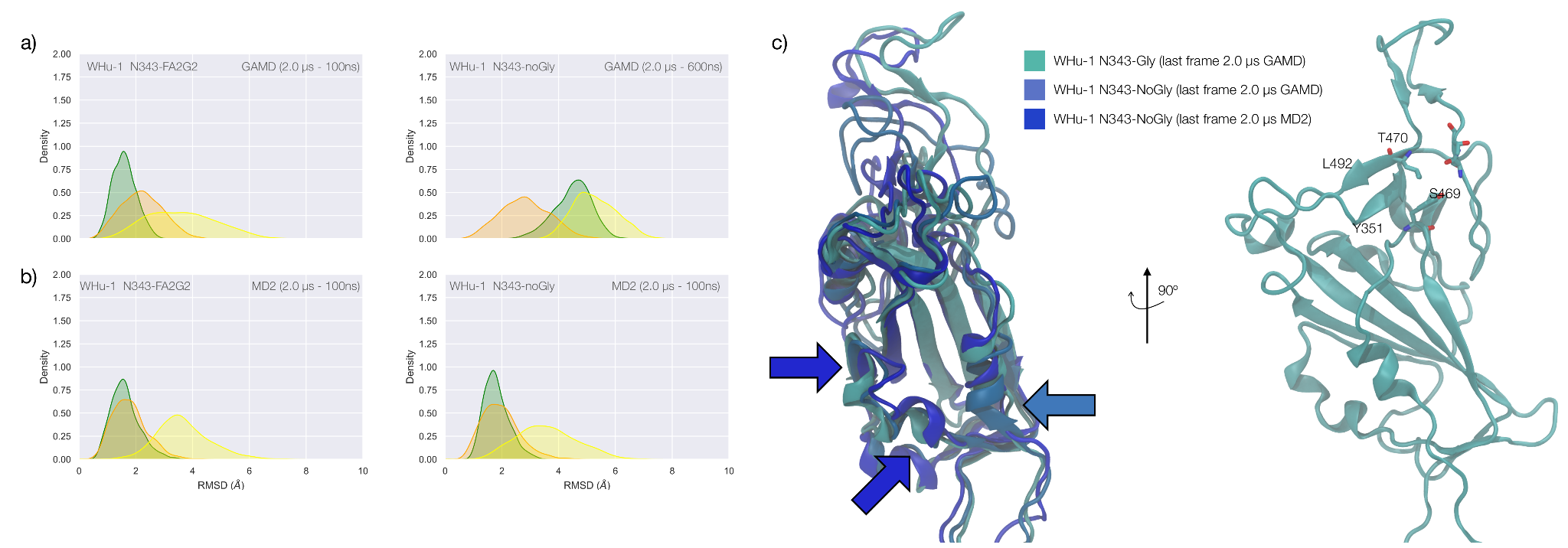

**Figure S.2 Panel a)** KDE plots of the backbone RMSD values calculated relative to frame 1 (t = 0) of the GaMD trajectory for Region 1 (green) aa 337-353, Region 2 (yellow) aa 439-506, and Region 3 (orange) aa 411-426 of the N343 glycosylated and non glycosylated WHu-1 RBD. The GaMD simulations were started from the structure of the RBD in PDB 6M0J. The first 100 ns of the N343 glycosylated RBD trajectory were considered part of the conformational equilibration and not included in the data analysis. The first 600 ns of the trajectory obtained for the N343 non glycosylated WHu-1 RBD were considered part of the conformational equilibration and not included in the data analysis. **Panel b)** KDE plots of the backbone RMSD values calculated relative to frame 1 (t = 0) of the conventional MD trajectory MD2 (see details above). MD2 was started from the conformation of the RBD in PDB 6M0J. The first 100 ns of both trajectories were considered part of the conformational equilibration and not included in the data analysis. **Panel c)** Graphical representation of the structural alignment of N343 glycosylated and non-glycosylated RBDs from the last frames of the GaMD and MD2 trajectories. Colour coded arrows (see legend) indicate where the main conformational changes leading to the tightening of the helices occur in each system. Proteins are represented by cartoons and N343 glycan is not represented for clarity. On the right-hand side a rotation of the RBD by 90^o^ clockwise shows how the hydrophilic loop in Region 2 is still well connected to Y351 in Region 1 at the end the GaMD N343 glycosylated WHu-1 RBD. Rendering done with VMD (<https://www.ks.uiuc.edu/Research/vmd/>) and KDE analysis with seaborn (<https://seaborn.pydata.org/>).

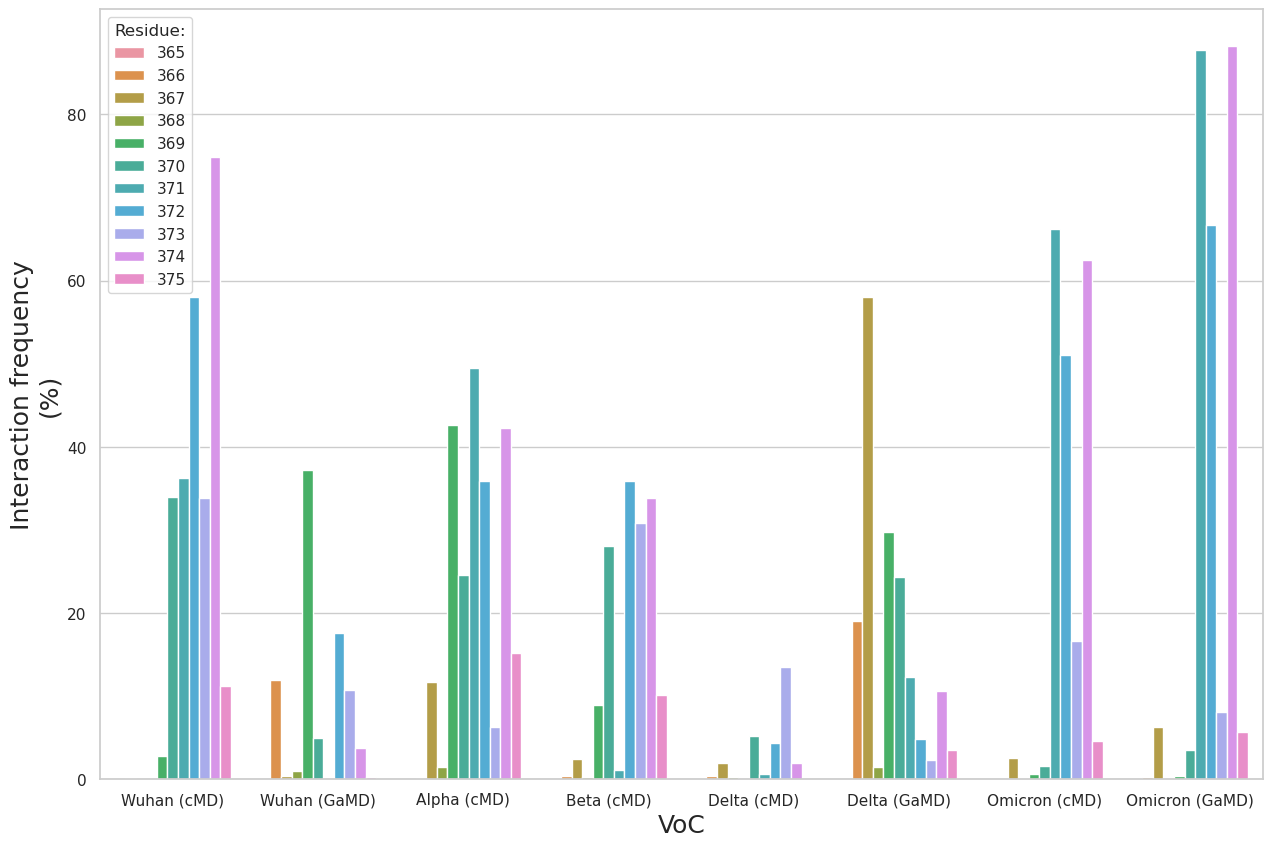

**Figure S.3** Bar plot of the interaction frequencies (%) of the N343 *N-*glycan with the different residues within the aa 365-375 loop for each VoC. The interactions include both hydrogen bonding and dispersion (van der Waals) contacts.

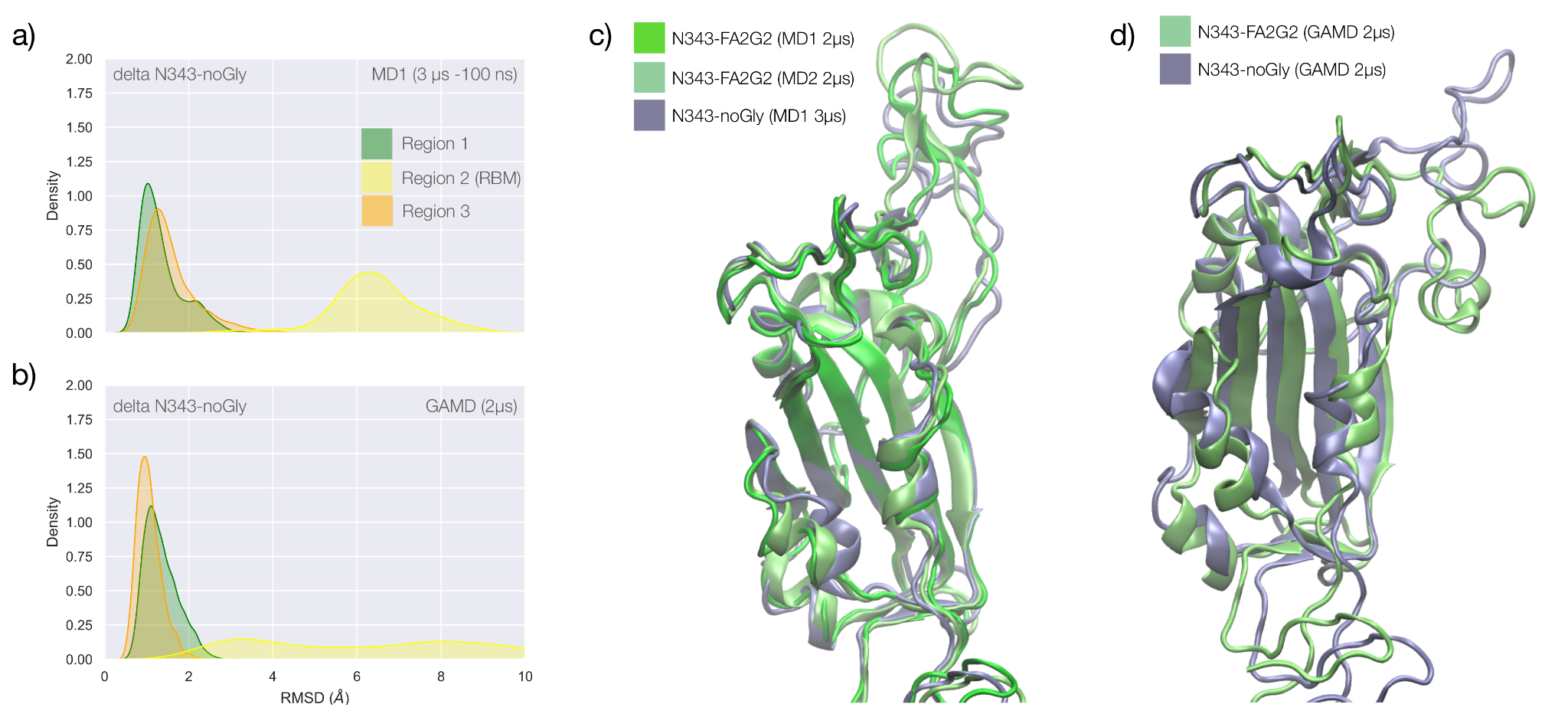

**Figure S.4 Panel a)** KDE plot of the backbone RMSD values calculated relative to frame 1 (t = 0) of the MD1 trajectory for Region 1 (green) aa 337-353, Region 2 (yellow) aa 439-506, and Region 3 (orange) aa 411-426 of the N343 non-glycosylated delta RBD. **Panel a)** KDE plot of the backbone RMSD values calculated relative to frame 1 (t = 0) of the GaMD trajectory of the N343 non-glycosylated delta RBD. **Panel c)** Graphical representations of the delta RBD structures from the last frame of the conventional simulation MD1 (N343 glycosylated and non-glycosylated) and MD2 (N343 glycosylated) with colourings indicated in the legend. Protein is represented with cartoons and the N343 and N331 glycans are omitted for clarity. **Panel c)** Graphical representations of the structurally aligned delta RBD structures from the last frame of the accelerated simulation GaMD (N343 glycosylated and non-glycosylated). Protein is represented with cartoons and the N343 and N331 glycans are omitted for clarity. Rendering done with VMD (<https://www.ks.uiuc.edu/Research/vmd/>) and KDE analysis with seaborn (<https://seaborn.pydata.org/>).

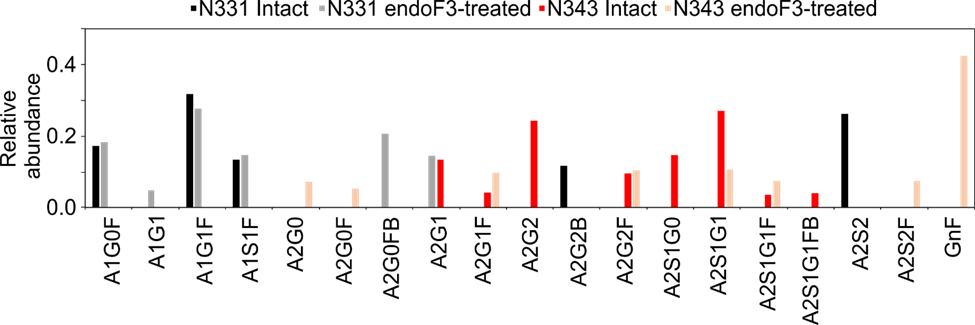

**Figure S.5.** Relative abundance of *N*-glycans at N331 and N343 on the WT RBD before and after endoF3 treatment.

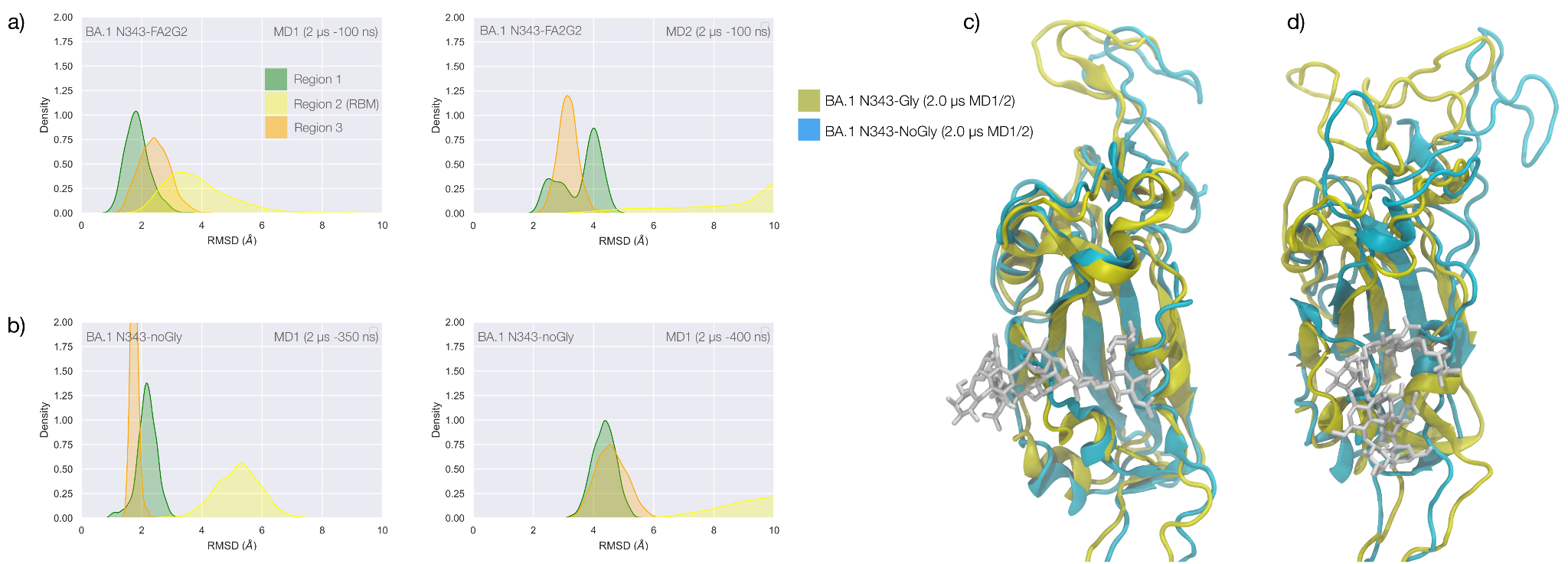

**Figure S.6. Panel a)** KDE plot of the backbone RMSD values calculated relative to frame 1 (t = 0) of the MD1 (left) and MD2 (right) trajectories for Region 1 (green) aa 337-353, Region 2 (yellow) aa 439-506, and Region 3 (orange) aa 411-426 of the glycosylated omicron (BA.1) RBD. **Panel b)** KDE plot of the backbone RMSD values calculated relative to frame 1 (t = 0) of the MD1 and MD2 trajectories (see details above) of the non-glycosylated omicron (BA.1) RBD. **Panel c)** Graphical representation of the structural alignment of the glycosylated (protein in yellow cartoons and N343-FA2G2 in white sticks, N331 omitted for clarity) and non-glycosylated (protein in cyan cartoons) of the omicron (BA.1) RBD from MD1. **Panel d)** Graphical representation of the structural alignment of the glycosylated (protein in yellow cartoons and N343-FA2G2 in white sticks, N331 omitted for clarity) and non-glycosylated (protein in cyan cartoons) of the omicron (BA.1) RBD from MD2. Structures correspond to the last frames of the trajectories, see details in the legend. Rendering done with VMD (<https://www.ks.uiuc.edu/Research/vmd/>) and KDE analysis with seaborn (<https://seaborn.pydata.org/>).

Casalino L, Gaieb Z, Goldsmith JA, Hjorth CK, Dommer AC, Harbison AM, Fogarty CA, Barros EP, Taylor BC, McLellan JS, Fadda E, Amaro RE. 2020. Beyond Shielding: The Roles of Glycans in the SARS-CoV-2 Spike Protein. *ACS Cent Sci* **6**:1722–1734. doi:10.1021/acscentsci.0c01056

D.A. Case, I.Y. Ben-Shalom, S.R. Brozell, D.S. Cerutti, T.E. Cheatham, III, V.W.D. Cruzeiro, T.A. Darden, R.E. Duke, D. Ghoreishi, M.K. Gilson, H. Gohlke, A.W. Goetz, D. Greene, R Harris, N. Homeyer, Y. Huang, S. Izadi, A. Kovalenko, T. Kurtzman, T.S. Lee, S. LeGrand, P. Li, C. Lin, J. Liu, T. Luchko, R. Luo, D.J. Mermelstein, K.M. Merz, Y. Miao, G. Monard, C. Nguyen, H. Nguyen, I. Omelyan, A. Onufriev, F. Pan, R. Qi, D.R. Roe, A. Roitberg, C. Sagui, S. Schott-Verdugo, J. Shen, C.L. Simmerling, J. Smith, R. SalomonFerrer, J. Swails, R.C. Walker, J. Wang, H. Wei, R.M. Wolf, X. Wu, L. Xiao, D.M. York and P.A. Kollman. 2018. AMBER.

Harbison AM, Fogarty CA, Phung TK, Satheesan A, Schulz BL, Fadda E. 2022. Fine-tuning the spike: role of the nature and topology of the glycan shield in the structure and dynamics of the SARS-CoV-2 S. *Chem Sci*.

Jorgensen WL, Chandrasekhar J, Madura JD, Impey RW, Klein ML. 1983. Comparison of simple potential functions for simulating liquid water. *J Chem Phys* **79**:926–935. doi:10.1063/1.445869

Kirschner KN, Yongye AB, Tschampel SM, González-Outeiriño J, Daniels CR, Foley BL, Woods RJ. 2008. GLYCAM06: a generalizable biomolecular force field. Carbohydrates. *J Comput Chem* **29**:622–655. doi:10.1002/jcc.20820

Kitova EN, El-Hawiet A, Schnier PD, Klassen JS. 2012. Reliable Determinations of Protein–Ligand Interactions by Direct ESI-MS Measurements. Are We There Yet? *J Am Soc Mass Spectrom* **23**:431–441. doi:10.1007/s13361-011-0311-9

Maier JA, Martinez C, Kasavajhala K, Wickstrom L, Hauser KE, Simmerling C. 2015. Ff14SB: Improving the accuracy of protein side chain and backbone parameters from ff99SB. *J Chem Theory Comput* **11**:3696–3713. doi:10.1021/acs.jctc.5b00255

Miao Y, Feher VA, McCammon JA. 2015. Gaussian Accelerated Molecular Dynamics: Unconstrained Enhanced Sampling and Free Energy Calculation. *J Chem Theory Comput* **11**:3584–3595. doi:10.1021/acs.jctc.5b00436

Nguyen L, McCord KA, Bui DT, Bouwman KM, Kitova EN, Elaish M, Kumawat D, Daskhan GC, Tomris I, Han L, Chopra P, Yang T-J, Willows SD, Mason AL, Mahal LK, Lowary TL, West LJ, Hsu S-TD, Hobman T, Tompkins SM, Boons G-J, de Vries RP, Macauley MS, Klassen JS. 2021. Sialic acid-containing glycolipids mediate binding and viral entry of SARS-CoV-2. *Nat Chem Biol* 1–10. doi:10.1038/s41589-021-00924-1

Wang J, Arantes PR, Bhattarai A, Hsu RV, Pawnikar S, Huang Y-MM, Palermo G, Miao Y. 2021. Gaussian accelerated molecular dynamics (GaMD): principles and applications. *Wiley Interdiscip Rev Comput Mol Sci* **11**. doi:10.1002/wcms.1521

Kitova, EN, El-Hawiet A, Schnier PD, Klassen JS. 2012. Reliable determinations of protein-ligand interactions by direct ESI-MS measurements. Are we there yet? *J Am Soc Mass Spectrom* **23**:431–441
